## Supplementary Figures for "Local Genetic Sex Differences in Quantitative Traits"

Affiliations:

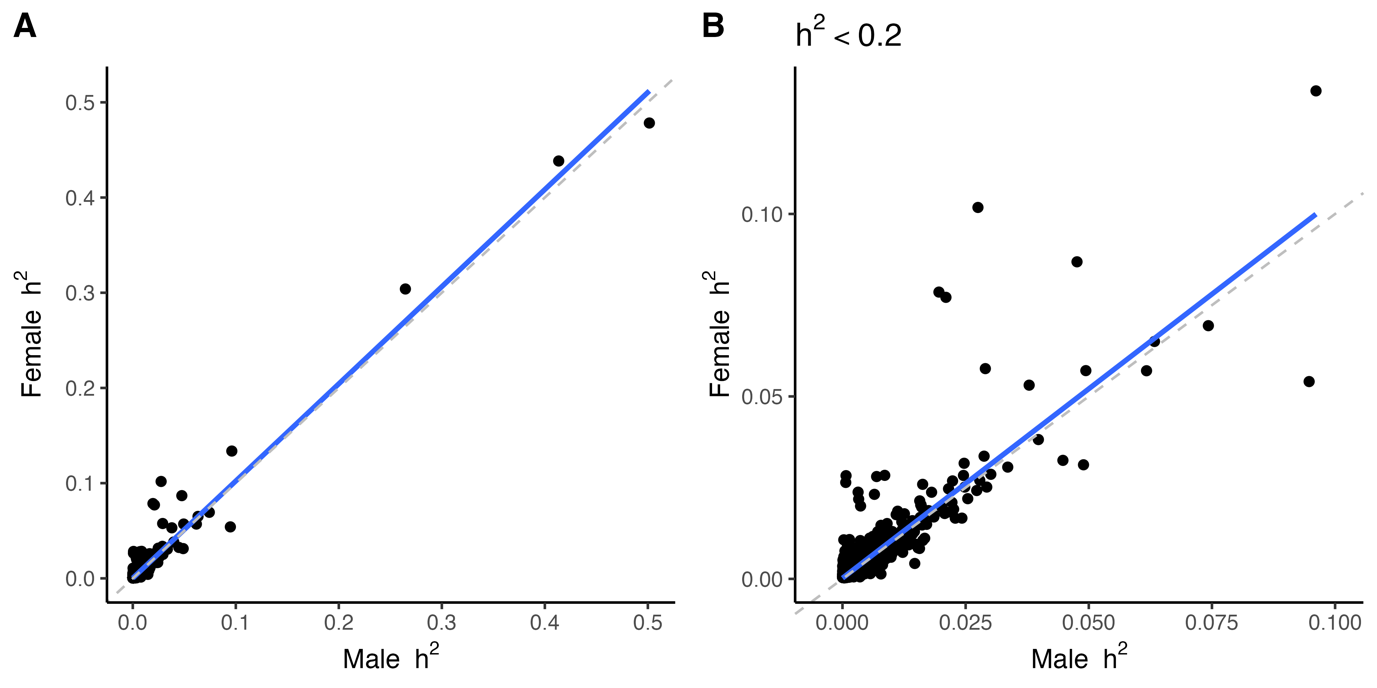

**Supplementary Figure 1**. Scatter plots of male and female local heritability (*h*^2^_local_) estimates. The left plot depicts all loci for all traits. The right plot depicts all loci with *h*^2^_local_ < 0.2. The three outlying *h*^2^_local_ in the left plot lead to an upward bias in the correlation estimate of male and female *h*^2^_local_s. Removal of these outliers reduces the correlation from *r*_pearson_ = 0.98 to *r*_pearson_ = 0.89. The grey dashed line shows the identity line. The regression line is depicted in blue.

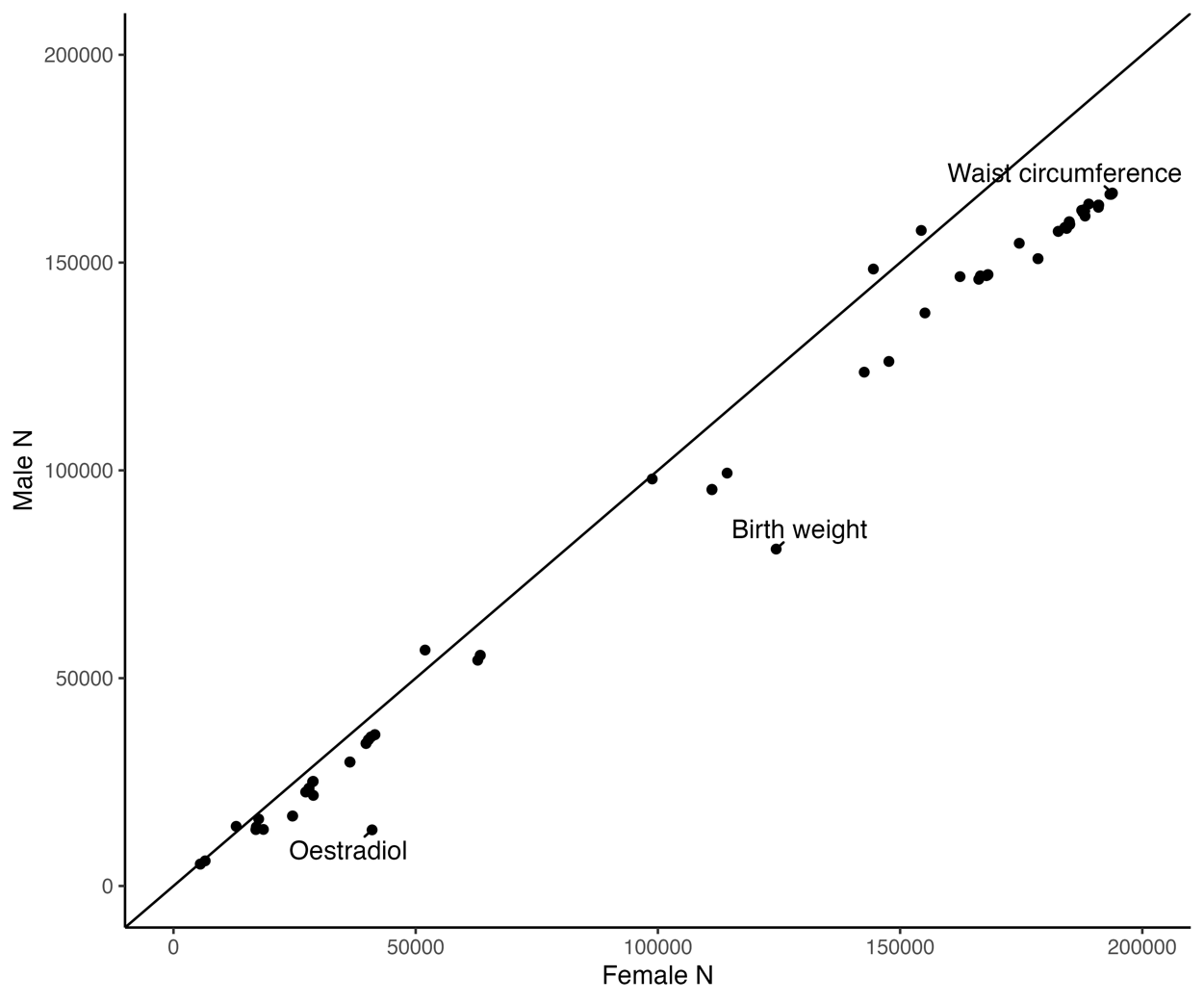

**Supplementary Figure 2**. Scatter plot of male and female GWAS sample sizes. The identity line (x=y) is depicted in black. Due to the UK Biobank female sampling bias, most traits have larger sample sizes for females. The median female-to-male sample size ratio is 1.16.

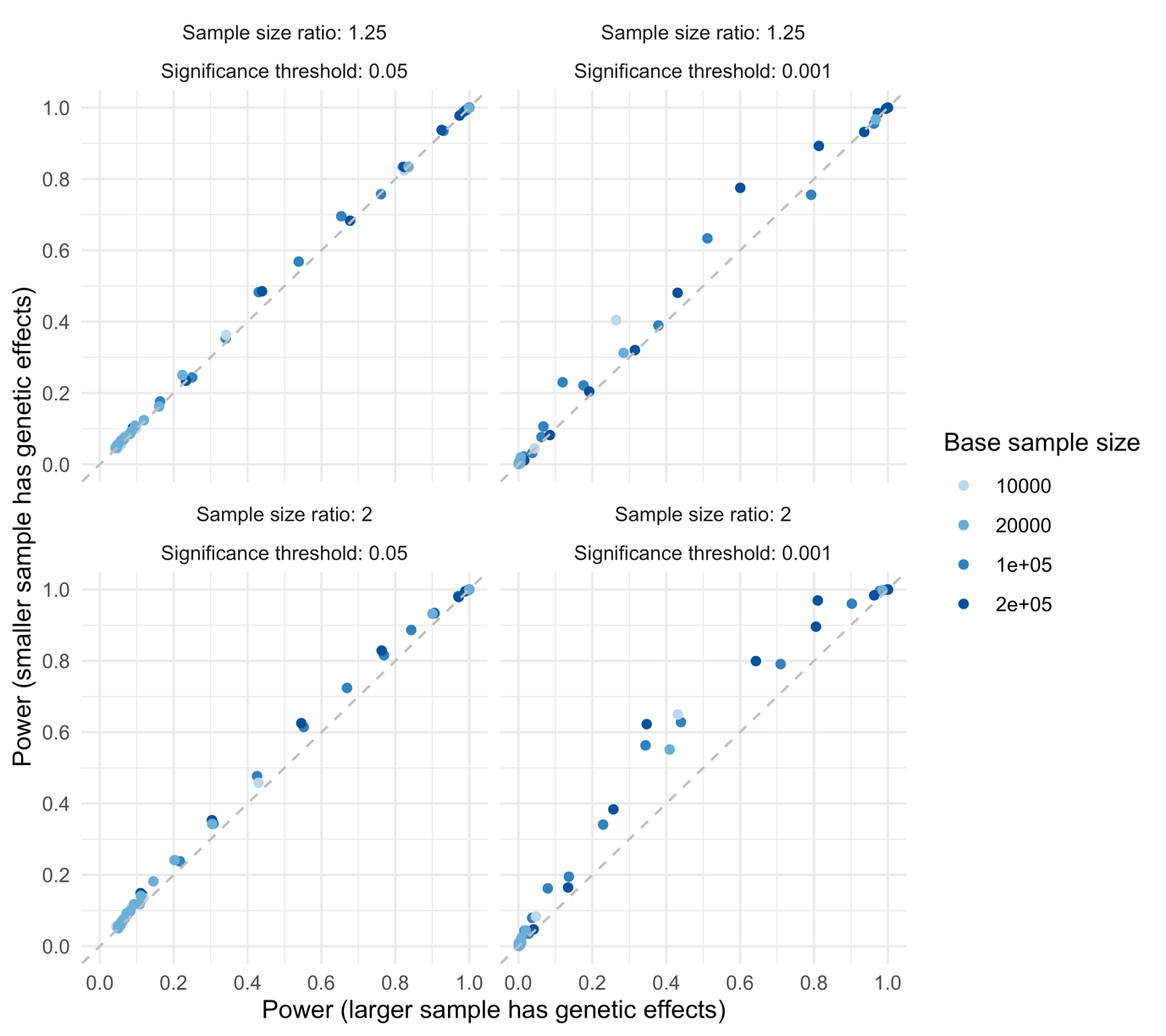

**Supplementary Figure 3**. Scatter plots of simulation results evaluating power to detect heritability differences when sample sizes differ between males and females. In each condition, samples of different sizes were created, and a phenotype was simulated with genetic effects in only one of the two samples (with no heritability in the other sample), and the power to detect this difference in heritability was computed. Two power values were computed for each condition, and are shown in the scatter plots: power to detect the difference in heritability with the genetic effects only in the larger sample (x-axis), and power with the genetic effects only in the smaller sample (y-axis). Sample sizes were set by selecting a base sample size for the smaller sample, then setting the larger sample at either 1.25 (top row) or 2 (bottom row) times that value. The heritability was varied from 0.01% to 1%, and power was evaluated at significance thresholds of 0.05 (left column) and 0.001 (right column). As shown, some asymmetry in power emerges at more pronounced differences in sample size, especially at lower significance thresholds, with more power if the higher heritability is in the smaller rather than the larger sample. See Methods for a detailed description of the simulations.

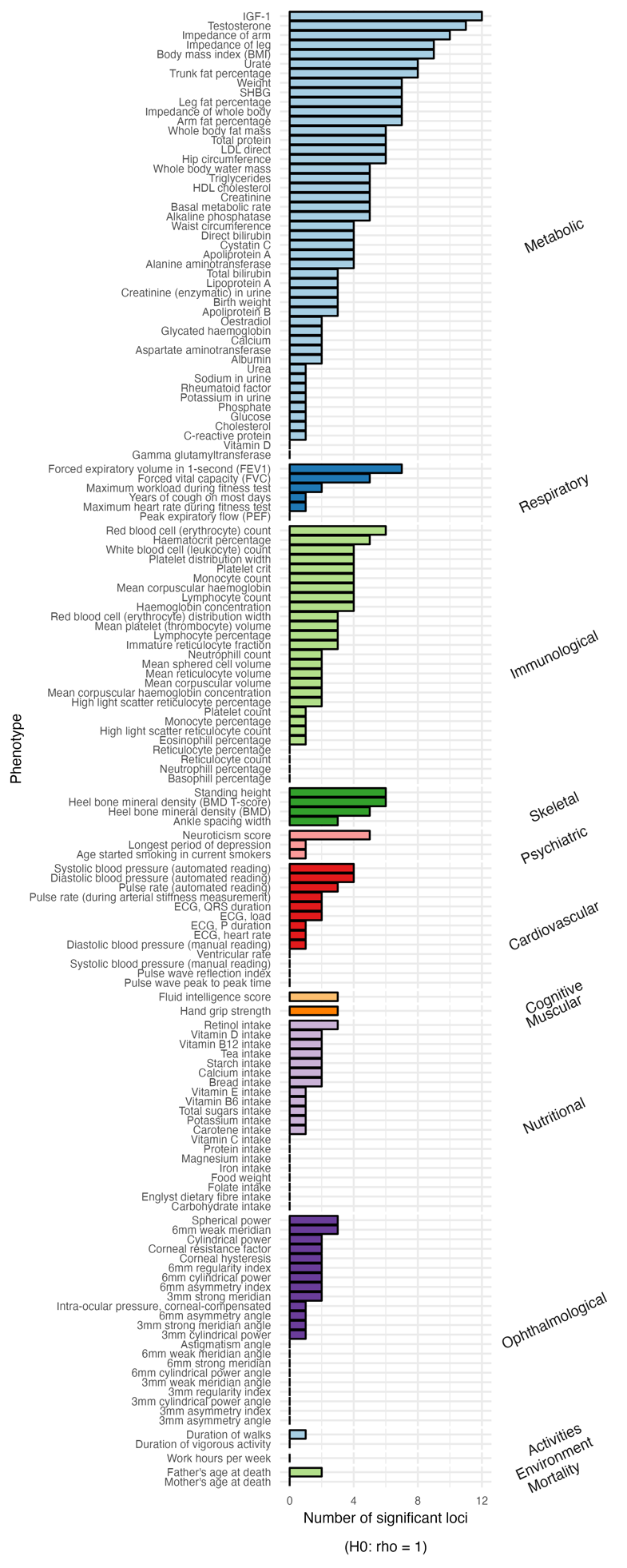

**Supplementary Figure 4.** Number of loci with local genetic correlations (*r*_g, local_) that are significantly different from one. We used a Bonferroni-corrected significance threshold of *p* < 0.05 / 11259 = 4.44e-06.

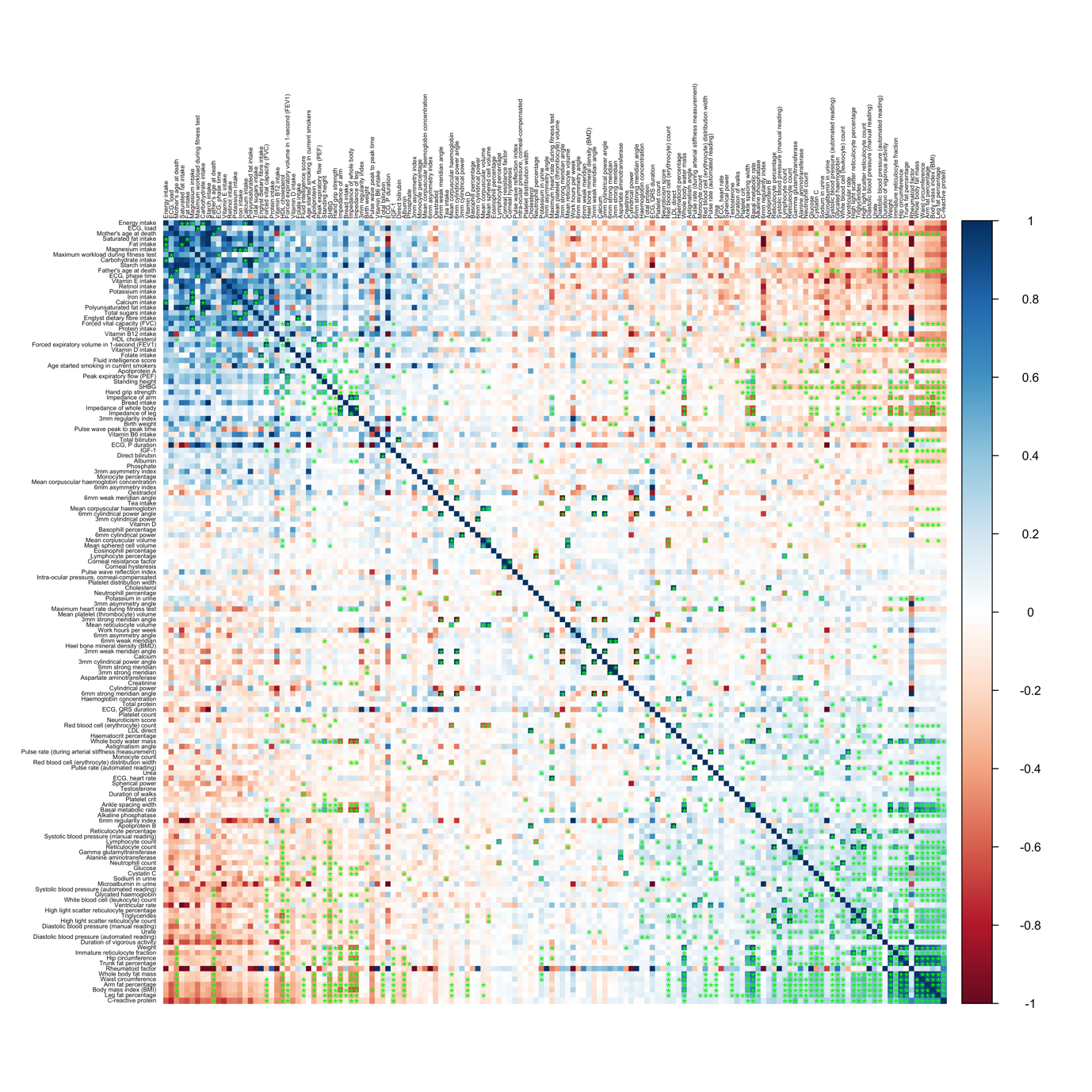

**Supplementary Figure 5**. Correlogram for global genetic correlations in females computed with LD Score Regression (LDSC). The plot was created with the corrplot v.0.92 R library. Traits were ordered based on the first principal component. LDSC p-values were Bonferroni-corrected for the total number of computed trait pairs across males and females (2.07e-06 = 0.05 / 24181) and significant correlations are marked with a green asterisk.

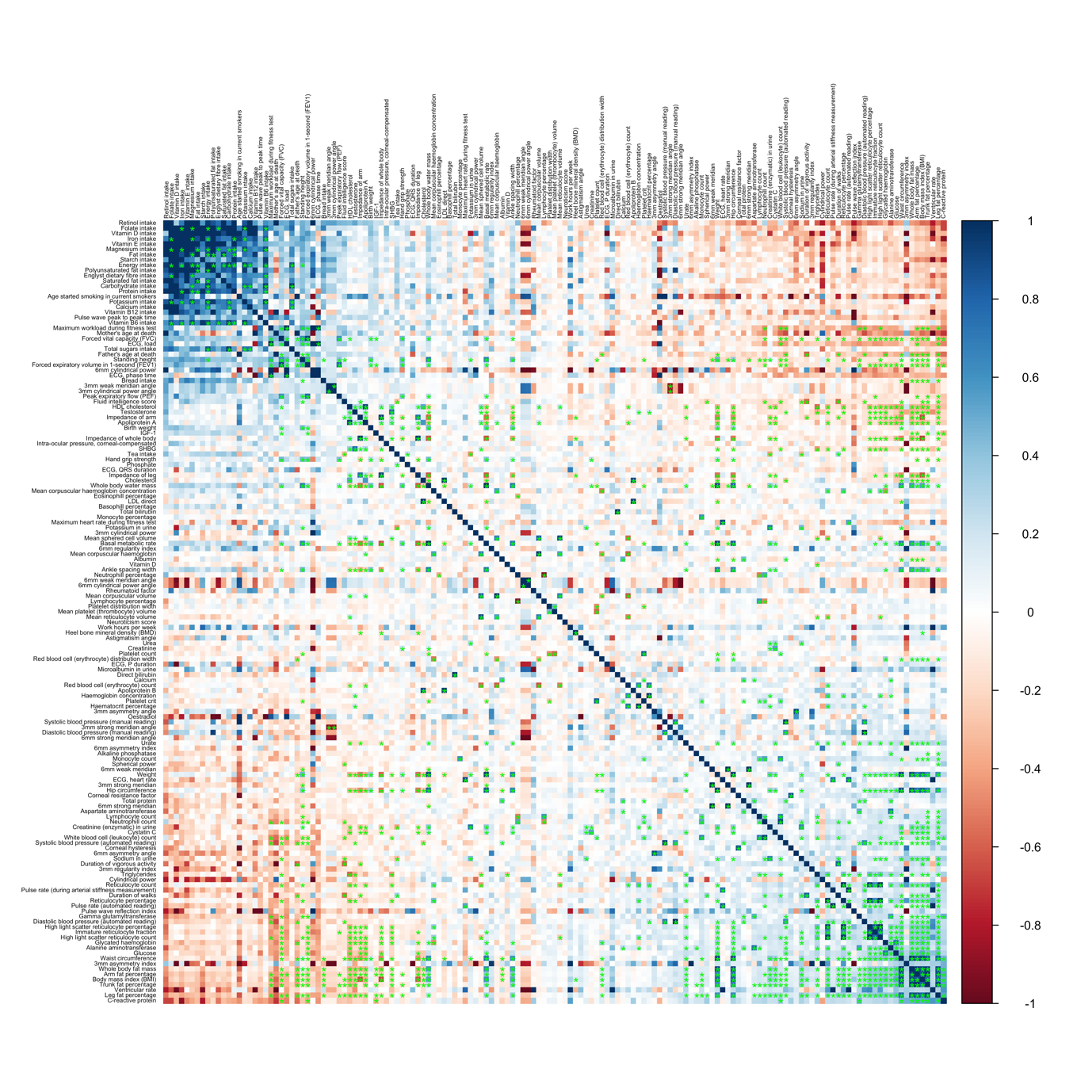

**Supplementary Figure 6**. Correlogram for global genetic correlations in males computed with LD Score Regression (LDSC). The plot was created with the corrplot v.0.92 R library. Traits were ordered based on the first principal component. LDSC p-values were Bonferroni-corrected for the total number of computed trait pairs across males and females (2.07e-06 = 0.05 / 24181) and significant correlations are marked with a green asterisk.

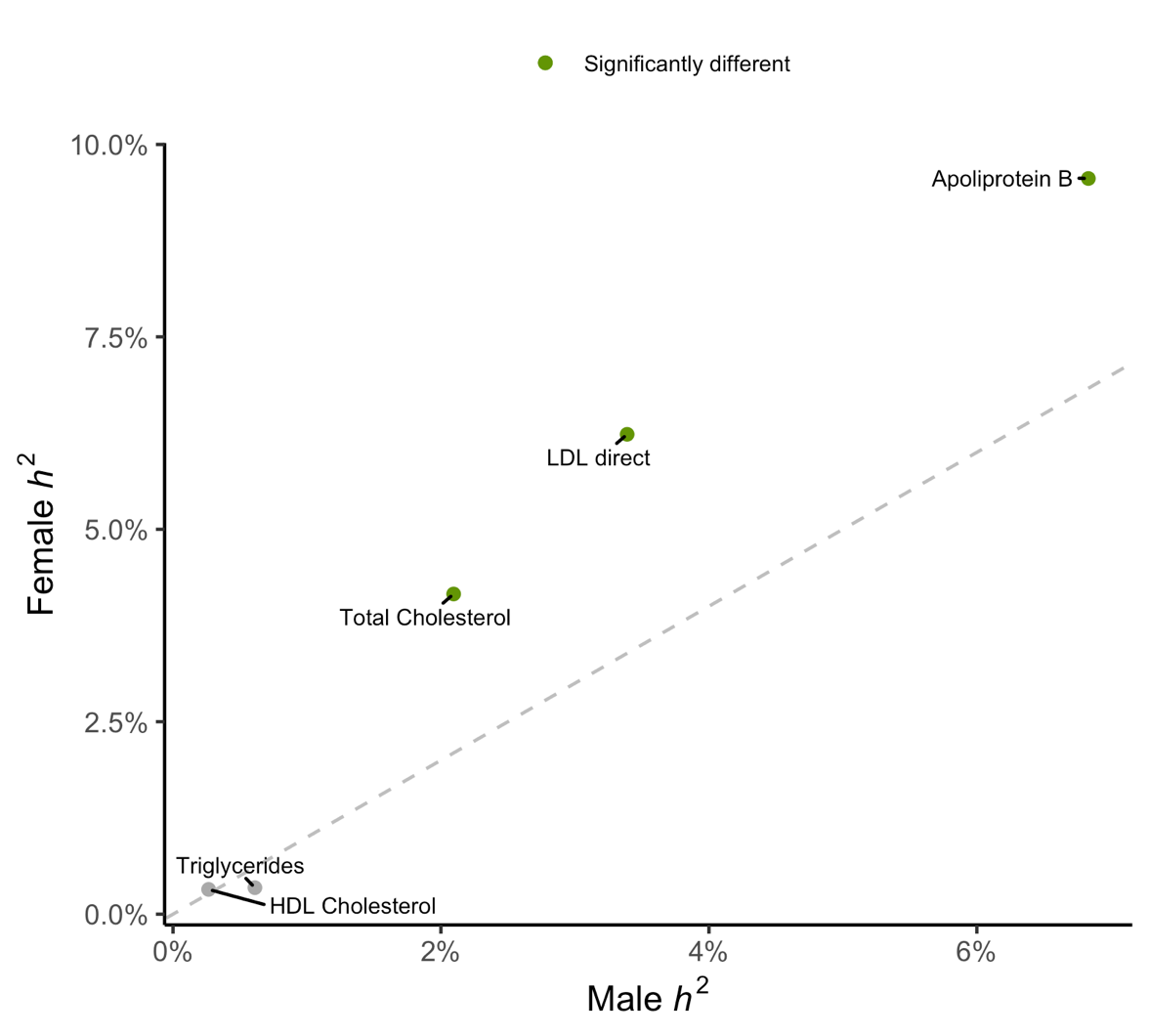

**Supplementary Figure 7**. Local heritability estimates for 2495 loci across five lipid-related traits in males and females. To test for significantly different heritability estimates, a Bonferroni-corrected threshold of *p* = 0.05 / (2495 x 157) = 1.28e-07 was used.
